# Cell-specific cargo delivery using synthetic bacterial spores

**DOI:** 10.1101/2020.02.13.947606

**Authors:** Minsuk Kong, Taylor B. Updegrove, Maira Alves Constantino, Devorah L. Gallardo, I-Lin Wu, David J. Fitzgerald, Kandice Tanner, Kumaran S. Ramamurthi

**Author notes:** To whom correspondence should be addressed. (D.J.F.) (K.T.), or; (K.S.R.).

## Abstract

SSHELs are synthetic bacterial spore-like particles wherein the spore’s cell surface is partially reconstituted around 1 µm-diameter silica beads coated with a lipid bilayer. Via a unique cysteine engineered in one of the surface proteins, the surface of SSHELs may be covalently decorated with molecules of interest. Here, we modified SSHELs with an affibody directed against HER2, a cell surface protein overexpressed in some breast and ovarian cancer cells, and loaded them with the chemotherapeutic agent doxorubicin. Drug-loaded SSHELs reduced tumor growth with lower toxicity in a mouse tumor xenograft model compared to free drug by specifically binding to HER2-positive cancer cells. We show that SSHELs bound to target cells are taken up and trafficked to acidic compartments, whereupon the cargo is released in a pH-dependent manner. Finally, we demonstrate that SHELLs can clear small tumor lesions in a complex tumor microenvironment in a zebrafish model of brain metastasis. We propose that SSHELs represent a versatile strategy for targeted drug delivery.

## INTRODUCTION

The efficacy of cancer chemotherapeutics can be abrogated by drug clearance and elimination, but also by delivery to non-specific sites, which can be dose-limiting. To diminish this problem, several methods employing nanometer-scale vehicles such as lipid vesicles have been devised to encase cargo for enhanced delivery to malignant cells within the tumor microenvironment ^1-3^. While these strategies can improve clearance and elimination, they typically rely on passive accumulation of the vehicle near solid tumors where accumulation is due to the abnormal vasculature ^4^. However, this effect can be unreliable ^5^, necessitating additional strategies that actively target the delivery of cargo to specific tissue types ^6-8^.

We have previously reported the assembly of synthetic bacterial spore-like particles termed “SSHELs” (<u>S</u>ynthetic <u>S</u>pore <u>H</u>usk-<u>E</u>ncased <u>L</u>ipid bilayers) ^9^. Spores of *Bacillus subtilis* are surrounded by a protein “coat” ^10^, built atop a basement layer containing two structural proteins ^11^: a membrane anchor SpoVM, ^12-16^ and a cytoskeletal protein SpoIVA ^17,18^ that hydrolyzes ATP to drive its static polymerization^19,20^. We recently reconstituted these proteins around spherical lipid bilayers supported by silica beads and demonstrated that, by employing a SpoIVA engineered to contain a single reactive Cys residue, SSHELs can covalently display thousands of copies of molecules of interest ^9,21^, thereby providing a useful platform for targeted drug delivery to cell surface markers, including those on cancer cells.

In many epithelial cancers, receptor tyrosine kinases (RTKs) are mutated, overexpressed, or non-functional compared to the normal phenotype ^22^. One such receptor, human epidermal growth factor receptor 2 (HER2), is usually present at copy numbers that facilitate normal cell proliferation and organ development of the ovary, lung, breast, liver, kidney, and central nervous tissue ^23,24^. However, in ∼25% of breast cancers it can be overexpressed, leading to aberrancies in growth and cell signaling ^25^. Monoclonal antibodies targeting HER2 have been effective in some breast cancer patients ^26^, but in ovarian cancers, the utility of HER2-overexpression as a prognostic factor has been unclear ^27-30^ and clinical trials with anti-HER2 agents have been largely unsuccessful in effective treatment. One possibility is that the levels of tissue penetrance are variable such that levels needed for drug efficacy are not realized in each patient. A second is that cues from the microenvironments of human breast and ovarian tissues reduce drug efficacy. We therefore sought to designed cargo-loaded SSHELs towards HER2-positive cells using an affibody that displays high affinity towards HER2 ^31^. Here, we report the assembly of 1 µm-diameter SSHELs built atop mesoporous silica beads loaded with the model cargo doxorubicin, an effective FDA-approved drug that promotes DNA double strand breaks, but whose systemic administration causes side effects like hair loss, bone marrow suppression, and vomiting ^32^. We demonstrate that doxorubicin-loaded SSHELs targeting HER2-positive cells effectively reduced tumor growth in a mouse tumor xenograft model of HER2-positive ovarian cancer. Growth retardation with doxorubicin-loaded SSHELs was superior to an equivalent dose of doxorubicin alone. Mechanistically, this was achieved by selective binding of particles to HER2-positive cells that were subsequently internalized and trafficked to acidified compartments, whereupon the drop in pH triggered cargo release. Finally, in a zebrafish model of breast cancer metastasis to the brain, SSHELs reduced tumor burden in this complex microenvironment. We propose that SSHELs may represent a versatile therapeutic vehicle whose targeting specificity and cargo are easily modified depending on the application.

## RESULTS

### Doxorubicin-encased SSHELs targeting HER2 inhibit tumor growth in a mouse xenograft model

To design biocompatible particles that can deliver cargo, we employed mesoporous silica beads containing 10 nm diameter pores and soaked them in a solution of doxorubicin (Fig. 1A). We chose doxorubicin as a model therapeutic agent due to its intrinsic red fluorescence that can be exploited to monitor drug loading and release kinetics, and because its p*K*_a_ of 8.2 allowed facile loading into the negatively charged mesoporous beads. We then applied a phospholipid bilayer composed of phosphatidylcholine to create spherical supported lipid bilayers (SSLBs) ^33^. Next, we applied synthesized SpoVM peptide to the SSLBs, after which we incubated the SpoVM-coated particles with purified SpoIVA in the presence of ATP to promote SpoIVA polymerization ^11^, resulting in stable SSHEL particles. The purified SpoIVA was a variant in which a native Cys residue had been substituted, and a single unpaired Cys residue was engineered into the N-terminus ^9^ to allow modification with *trans*-cyclooctene (TCO).

**Figure 1.**
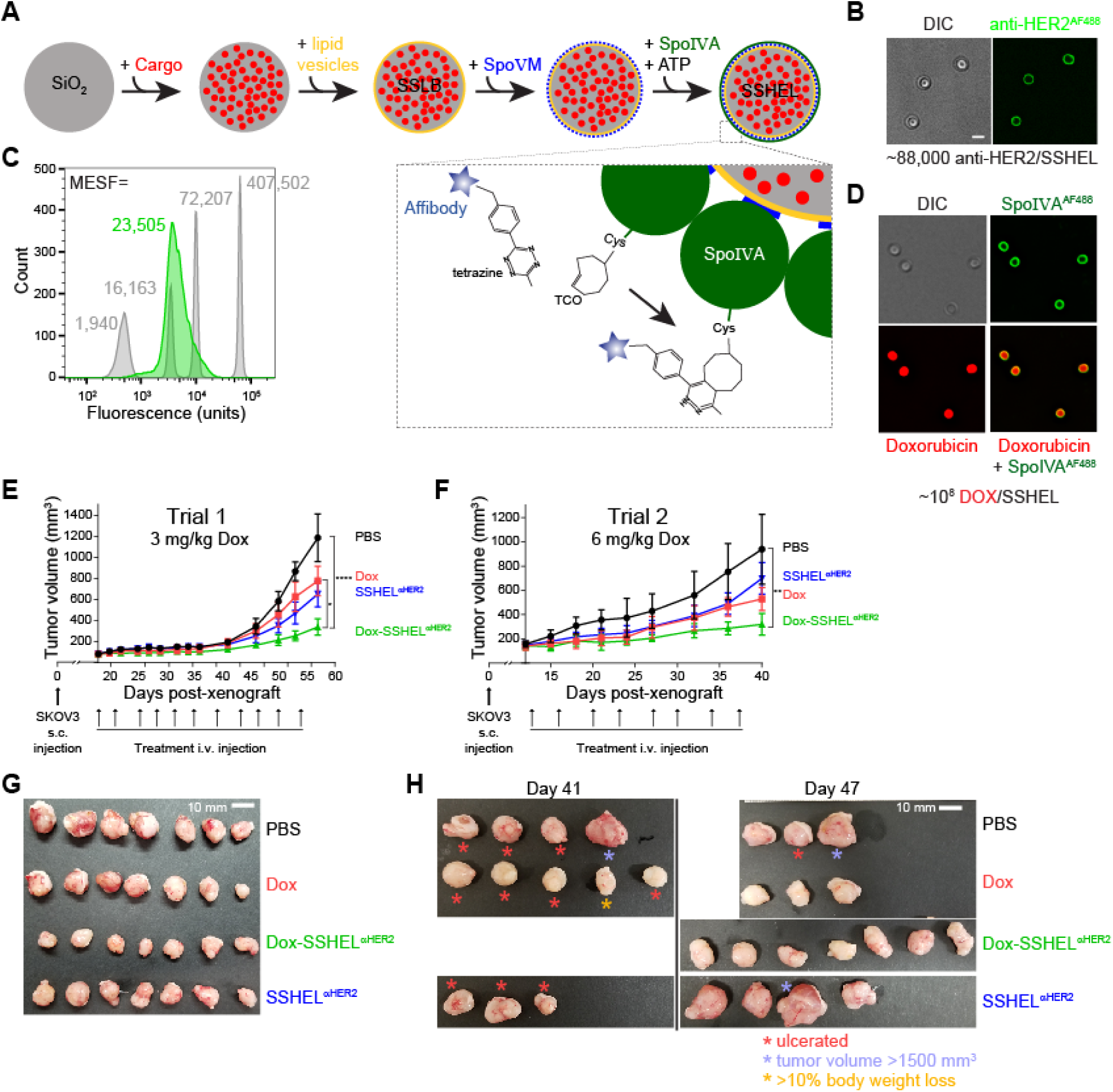
Doxorubicin delivery via SSHELs targeting HER2-positive cells reduces tumor burden in a mouse tumor xenograft model. (A) Schematic representation of SSHEL particle assembly. 1 µm-diameter mesoporous silica beads (gray, SiO_2_) are loaded with cargo (doxorubicin, red), after which a lipid bilayer (phosphatidylcholine) is applied to the surface (yellow) to create cargo-encased spherical supported lipid bilayers (SSLB). SSLBs are then incubated with SpoVM peptide (blue) and SpoIVA protein (green) and ATP to promote SpoIVA polymerization. Inset: SpoIVA contains an engineered Cys conjugated to trans-cylcooctene (TCO). Incubation with anti-HER2 affibody (blue star) conjugated to cognate click chemistry molecule tetrazine drives formation of a covalent dihydropyridazine bond that results in affibody display on the SSHEL surface. (B) Fluorescence micrograph of SSHELs displaying covalently-linked affibodies labeled with Alexa Fluor 488 (AF488) fluorescent dye. Left: SSHELs visualized using DIC; right: fluorescence from AF488. (C) Fluorescence from SSHELs displaying anti-HER2^AF488^ (green) measured using flow cytometry and compared to fluorescence from beads displaying known quantity of molecules of equivalent soluble fluorochrome (MESF) to calculate the number of anti-HER2^AF488^ displayed per SSHEL particle. (D) Fluorescence micrographs of doxorubicin-loaded SSHELs made with SpoIVA^AF488^. Top left: DIC; top right: fluorescence from SpoIVA^AF488^; bottom left: fluorescence from doxorubicin; bottom right: overlay, doxorubicin and SpoIVA^AF488^. Scale bars in B and D: 1 µm. (E-F) Athymic nude mice were inoculated subcutaneously (s.c.) with SKOV3 HER2-positive ovarian cancer cells. When tumor volume reached ∼100 mm^3^, mice were treated intravenously (i.v.) with PBS (black circles), (E) 60 µg or (F) 120 µg doxorubicin (red squares), doxorubicin-loaded SSHELs containing an equivalent dose of doxorubicin (green triangles), or an equivalent number of SSHELs without cargo (blue inverted triangles) at the days post xenograft indicated with arrows(18, 21, 25, 28, 32, 35, 39, 43, 46, 50, 54 for trial 1; 13, 16, 20, 23, 27, 30, 34, 37 for trial 2) and tumor volumes were measured. Data points represent mean values; errors are S.D.; n=7 mice. P-values: *<.05; ****<.001. (G-H) Tumors excised from mice in (E-F), respectively, at (G) day 60, or (H) day 41 (H, left) or day 47 (H, right). Red asterisk: ulcerated tumor; blue asterisk: tumors >1500 mm^3^; orange asterisk: tumor excised from mouse exhibiting >10% body weight loss. Scale bars: 10 mm.

As a model target cell, we chose a set of well-studied “HER2-positive” breast cancer cells. We therefore covalently attached to the surface of TCO-modified SSHELs a fluorescently-labeled affibody that binds to HER2 with a high affinity (*k*_d_ = 22 pM) that had been conjugated, via an engineered C-terminal Cys, with tetrazine, which undergoes a copper-free inverse-electron-demand Diels-Alder reaction with TCO (Fig. 1A) ^34,35^. Fluorescence microscopy revealed that the labeled affibodies uniformly coated the SSHELs (Fig. 1B). Quantitative flow cytometry indicated that each SSHEL particle displayed a mean of 23,500 ± 2,400 (n=3 independent labeling experiments with ∼10,000 particles) molecules of equivalent soluble fluorophore (MESF) (Fig. 1C) which, after accounting for labeling efficiency translated to each particle displaying 94,000 ± 9,600 anti-HER2 affibodies. Fluorescence microscopy of doxorubicin-loaded SSHELs (DOX-SSHEL) constructed using fluorescently labeled SpoIVA indicated that doxorubicin was successfully retained in the center of the particles after SpoIVA polymerized around the periphery of the particles (Fig. 1D). Extraction of encased doxorubicin with detergent and measurement of its fluorescence intensity revealed that each particle contained 1.5 × 10^8^ ± 1.4 × 10^7^ doxorubicin molecules.

For in vivo testing, athymic nude mice were inoculated subcutaneously in the flank with SKOV3 cells, a HER2-positive ovarian cancer cell line. When the tumors reached ∼100 mm^3^, mice were treated intermittently eleven times intravenously with either PBS, free doxorubicin (3 mg/kg), DOX-SSHELs displaying the αHER2 affibody (DOX-SSHEL^αHER2^, using an equivalent doxorubicin dose), or empty SSHEL^αHER2^. In the PBS-treated mice, the mean tumor size (n=7) reached ∼1200 mm^3^ in 57 days, whereas treatment with doxorubicin reduced the mean tumor size to ∼800 mm^3^ by day 57 (Fig. 1E, G). Treatment with SSHEL^αHER2^, without drug, had a similar effect on mean tumor size as doxorubicin. However, in mice treated with DOX-SSHEL^αHER2^ mean tumor size was only ∼300 mm^3^. At this concentration of doxorubicin, we did not observe any appreciable signs of toxicity or body weight loss (Fig. S1A) in any treatment group. We then repeated the experiment using 6 mg/kg of doxorubicin. In this second experiment, tumors grew slightly faster, with mean tumor size (n=7) in PBS-treated mice reaching ∼1000 mm^3^ in 40 days (Fig. 1F). Additionally, several tumors displayed ulceration (4/7) and some tumors were larger than 1500 mm^3^ (2/7; Fig. 1H). As seen in the first trial, treatment with doxorubicin or SSHEL^αHER2^ reduced mean tumor size to ∼600 mm^3^ by day 40, but several tumors from doxorubicin-treated mice were ulcerated (4/7) and one mouse experienced >10% loss in body weight. Additionally, mice in this treatment group displayed pale skin (5/7) and exhibited tissue damage at the site of injection (3/7). In mice treated with SSHEL^αHER2^, we observed ulcerated tumors (3/7), and one tumor that was larger than 1500 mm^3^. In contrast, treatment with DOX-SSHEL^αHER2^ resulted in mean tumor size of ∼300 mm^3^ at day 40. Furthermore, none of the mice displayed significant weight loss (Fig. S1B) and none of the tumors from this treatment displayed ulceration (Fig. 1H). Administration of doxorubicin via SSHEL^αHER2^ therefore enhanced therapeutic efficacy while reducing the adverse effects of the drug.

### HER2^+^ cells specifically bind to and internalize SSHEL^αHER2^ particles

It was proposed that antibody treatment prevents shedding of the extracellular domain of HER2 and directs HER2 for internal degradation. To compare the mechanism for the increased efficacy of SSHEL-delivered drugs, we first tested the binding of fluorescently-labeled SSHEL^αHER2^ in vitro to two HER2^+^ cell lines: SKOV3 cells used in the mouse xenograft model (Fig. 1) and SKBR3 cells (breast cancer), and compared it to binding to two HER2^-^ breast cancer cell lines: MDA-MB-231 and MCF7. Particles were incubated at different multiplicities of incubation (MOI) with detached cells and analyzed using flow cytometry. With increasing MOI, SKBR3 and SKOV3 cells displayed increased fluorescence (Fig. 2A-B), whereas incubation of SSHEL^αHER2^ to MDA-MB-231 and MCF7 did not result in similar increases (Fig. 2C-D), suggesting preferential binding to HER2^+^ cells. To test if binding was mediated by HER2, we first incubated the cells with a 1000-fold excess of unlabeled anti-HER2 affibodies, then repeated the binding experiment with fluorescently labeled SSHEL^αHER2^. Preincubation with anti-HER2 affibodies blocked the binding of SSHEL^αHER2^ at MOI=100 (Fig. 2E-F), indicating that the binding of the particles was mediated by HER2 and was saturable. The specific binding of the SSHEL^αHER2^ particles to the surface of undetached HER2^+^ (Fig. 2G-H), but not HER2^-^ (Fig. 2I) cells was qualitatively confirmed using confocal microscopy.

**Figure 2.**
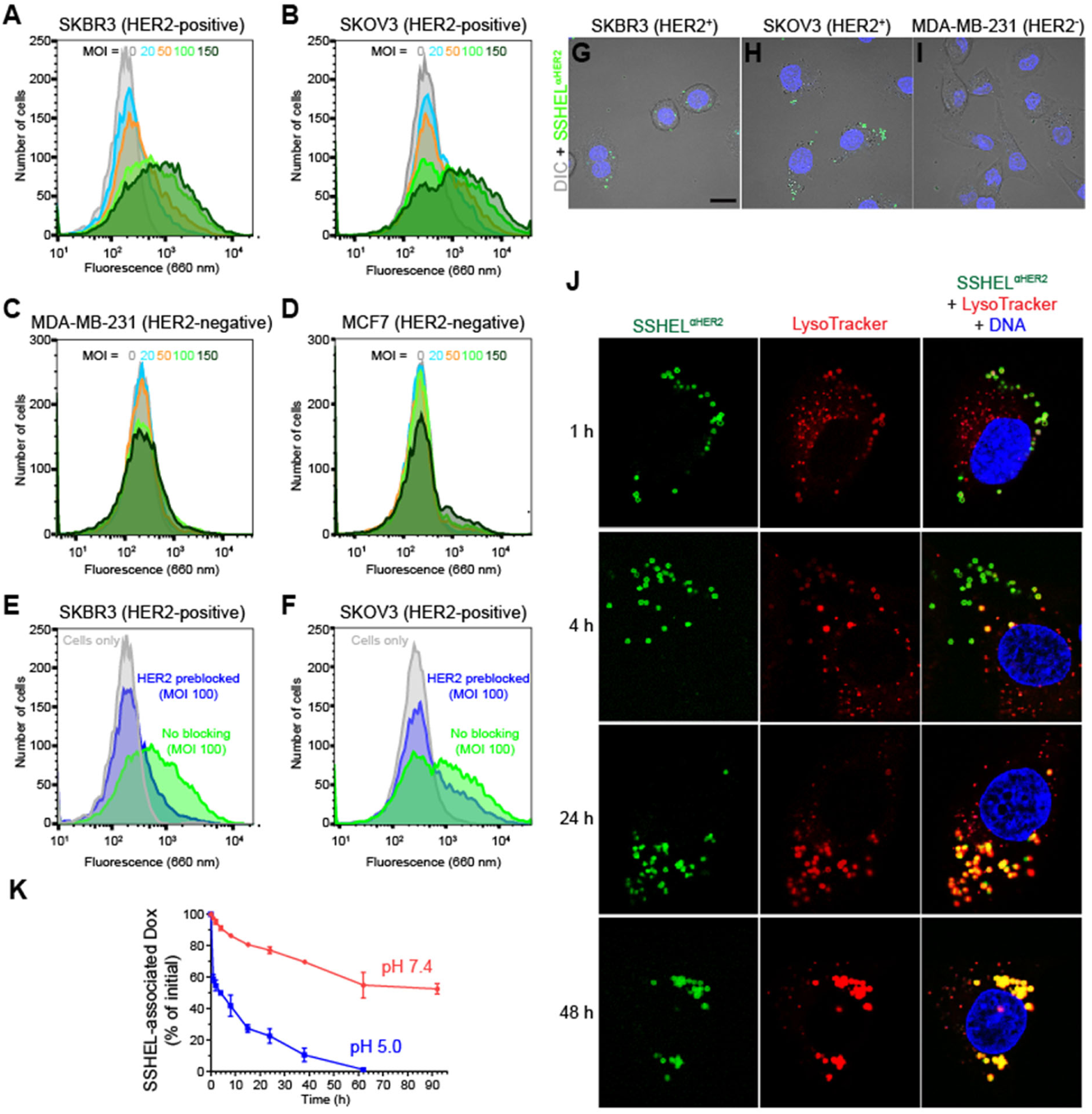
SSHELs displaying anti-HER2 bind specifically to HER2-positive cells and are trafficked to acidic compartments. (A-D) Fluorescently-labeled SSHEL^αHER2^ particles were incubated at different multiplicities of incubation (MOI) with HER2-positive (A-B; SKBR3 breast cancer (A) or SKOV3 ovarian cancer (B)) or HER2-negative (C-D; MDA-MB-231 (C) or MCF7 (D)) cells and binding was assessed using flow cytometry. (E-F) SKBR3 (E) or SKOV3 (F) cells were pre-incubated with an excess of anti-HER2 affibody, after which the pre-blocked cells were incubated with SSHEL^αHER2^ particles at MOI=100 and binding was assessed using flow cytometry. (G-I) Fluorescence micrographs of (G) SKBR3, (H) SKOV3, or MDA-MB-231 cells incubated with AF488-labeled SSHEL^αHER2^ particles. Shown is an overlay of DIC (gray) and fluorescence from AF488 (green). Scale bar: 20 µm. (J) Confocal micrograph of SKBR3 cells labeled with LysoTracker and DAPI (to visualize acidic compartments and DNA, respectively) incubated with AF488-labeled SSHEL^αHER2^ particles and visualized at the indicated time points. Left column: fluorescence from AF488 (green); center column: fluorescence from LysoTracker (red); right column: overlay of AF488, LysoTracker, and DAPI (blue); yellow indicates colocalization of AF488 and LysoTracker. (K) Release of doxorubicin from SSHELs over time (detected using intrinsic fluorescence of doxorubicin) over time reported as a percentage of initial amount of encapsulated doxorubicin when incubated in pH=7.4 (red) or pH=5.0 (blue). Data points are mean values; errors are S.D. (n=3).

We next incubated SKOV3 cells with AF488-labeled SSHEL^αHER2^ and monitored for 48 h the co-localization of particles with acidic organelles that were visualized using the fluorescent dye LysoTracker Red using confocal microscopy. Within 1 h of incubation, SSHEL^αHER2^ particles were located at the periphery of SKOV3 cells (Fig. 2J). Beginning at 4 h, SSHEL^αHER2^ particles began to co-localize with the LysoTracker Red dye, suggesting internalization of some of the particles and trafficking to acidic compartments. The extent of internalization increased over time, and by 48 h, the majority of cell associated SSHEL^αHER2^ particles colocalized with acidic compartments, consistent with the relatively slow trafficking of SSHEL^αHER2^ into acidic organelles.

We next tested if exposure of DOX-SSHELs to acidic pH could liberate the cargo by incubating the particles in either simulated serum (phosphate-buffered saline with 10% serum at pH 7.4) or simulated endosomal/lysosomal conditions (0.1 M citrate buffer at pH 5.0) and monitoring doxorubicin release over time. In acidic pH, encased doxorubicin was released with a half-life of 4 h, and nearly all the encased drug was released by 60 h (Fig. 2K). In contrast, in near-neutral pH, the release of doxorubicin was reduced, with more than 50% of the drug remaining associated within the particles even after 92 h, suggesting that SSHELs are largely stable under physiological conditions, but can be disrupted or destabilized in acidic conditions, such as those found upon trafficking to an acidified compartment.

### DOX-SSHEL^αHER2^ particles specifically reduce viability of HER2^+^ cells by inducing apoptosis

We next measured viability of different cell lines after incubation with DOX-SSHEL^αHER2^ at different MOIs. Incubation of the two HER2^+^ cell lines with DOX-SSHEL^αHER2^ resulted in the dose-dependent reduction of cell viability, whereas the viability of two HER2^-^ cell lines were either unaffected (MCF7) or reduced to only ∼65% at MOI=300 (MDA-MB-231; Fig. 3A). These MOIs corresponded to a half maximal inhibitory concentration (IC_50_) of encapsulated doxorubicin of 41 nM (SKBR3) and 86 nM (SKOV3), which was comparable to the IC_50_ of free doxorubicin for both cell lines (56 nM for SKBR3; 73 nM for SKOV3; Fig. 3B). As opposed to encapsulated doxorubicin, free doxorubicin reduced the viability of HER2^-^ cells and displayed an IC_50_ of 96 nM for MDA-MB-231 and 135 nM MCF7 cells. In contrast, unadorned SSHELs or the anti-HER2 affibody alone did not reduce the viability of any of the cell lines (Fig. 3C-D). Incubating cells with doxorubicin-loaded SSHELs displaying bovine serum albumin (DOX-SSHEL^BSA^). showed reduced, but non-discriminate, cytotoxicity against all cell lines tested (Fig. 3E), with an IC_50_ value approximately 3.3-fold (SKBR3) or 2.3-fold (SKOV3) higher than that of DOX-SSHEL^αHER2^, indicating the necessity of HER2-targeting for efficient and cell-selective cytotoxicity. Lastly, to verify that encapsulated cargo is responsible for cell toxicity, we incubated cells with SSHEL^αHER2^, without encapsulated doxorubicin. Although SSHEL^αHER2^ caused some reduction in viability of HER2^+^ cells at the highest MOI (Fig. 3F), its effect was much lower than that of DOX-SSHEL^αHER2^, suggesting that the combined action of encapsulated doxorubicin directed by the particles via the anti-HER2 affibody, was required to exert full anti-proliferative activity towards HER2^+^ cells.

**Figure 3.**
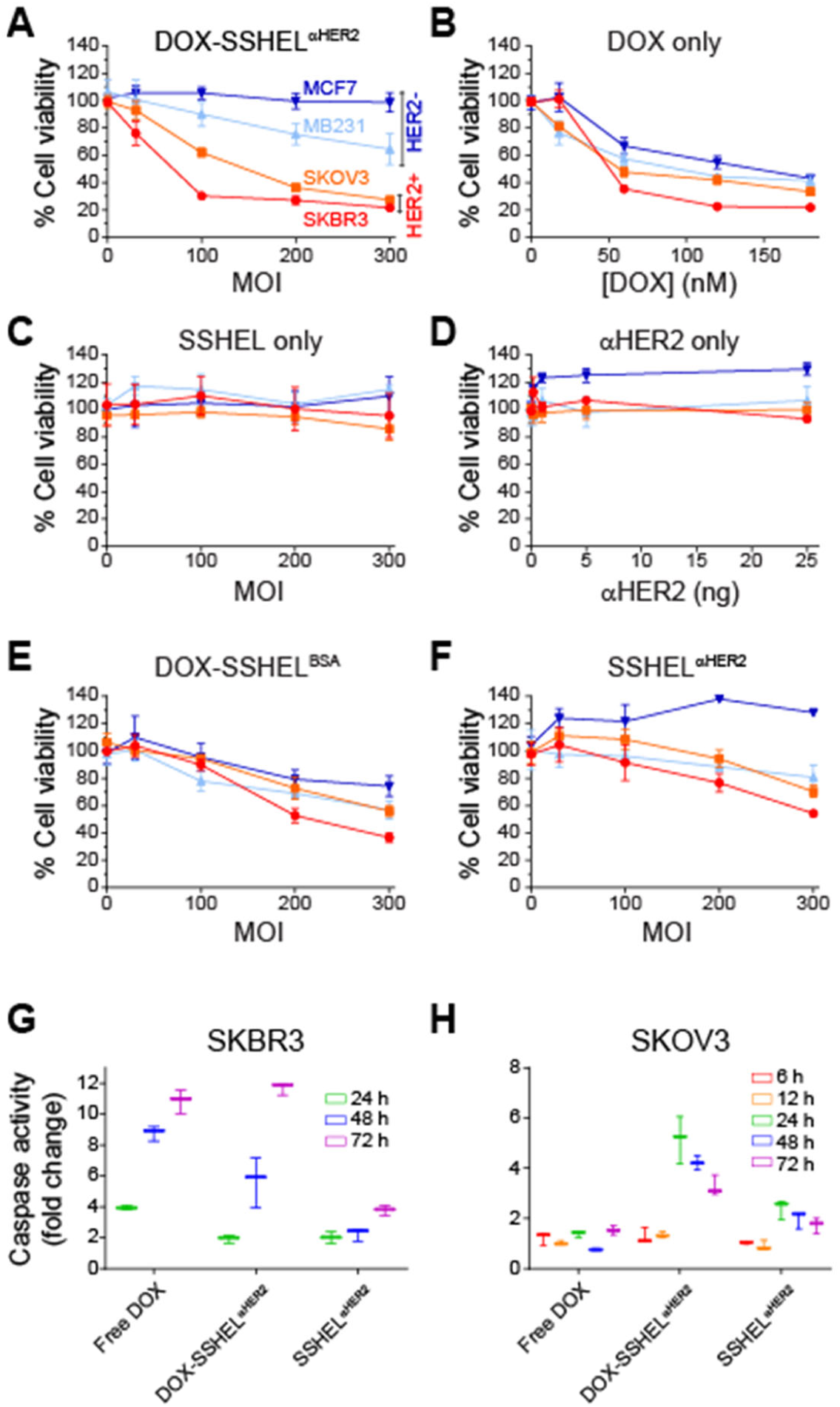
Doxorubicin-loaded SSHEL^αHER2^ specifically kills HER2-positive cells in vitro by inducing apoptosis. Cell viability of HER2-positive (SKBR3, red; SKOV3, orange) and -negative (MCF7, blue; MDA-MB-231, light blue) cells incubated for 96 h with (A) doxorubicin-loaded SSHEL^αHER2^ at different MOIs; (B) free doxorubicin at equivalent concentrations to that used in (A); (C) SSHEL without loaded doxorubicin or conjugation with anti-HER2 affibody; (D) anti-HER2 affibody only at equivalent quantity as present in (A); (E) doxorubicin-loaded SSHELs decorated with bovine serum albumin (BSA); or (F) SSHEL^αHER2^ without doxorubicin. Data points are mean values; errors are S.D. (n=3 independent experiments). (G-H) Caspase 3 and 7 activities of HER2-positive cancer cells (G) SKBR3 and (H) SKOV3 were measured at indicated time points after treatment with free doxorubicin (120 nM), unloaded SSHEL^αHER2^ (MOI=200), or doxorubicin-loaded-SSHEL^αHER2^ (MOI=200, equivalent to 120 nM doxorubicin). Data represent the average fold change compared to PBS-treated cells; errors are S.D. (n=3).

Treatment with doxorubicin has been reported to induce apoptosis in various tumor cell lines^36^. We therefore monitored induction of apoptosis by measuring the caspase 3 and 7 activity of HER2^+^ cells at different time points after treatment with DOX-SSHEL^αHER2^, unloaded SSHEL^αHER2^ (each at MOI= 200), or an equivalent dose of doxorubicin (120 nM). In contrast to unloaded SSHEL^αHER2^, doxorubicin and DOX-SSHEL^αHER2^ induced caspase activity of SKBR3 cells over time (Fig. 3G), albeit with differing kinetics (compare caspase induction at 48 h in Fig. 3G), suggesting that the cellular uptake rates between free drugs and SSHEL-encapsulated drugs may differ. For SKOV3 cells, 120 nM doxorubicin did not significantly induce caspase activity, even after 72 h (Fig. 3H), and increasing the concentration to 10 µM only resulted in an ∼two-fold increase (Fig S2), suggesting that the reduced viability of SKOV3 cells by doxorubicin shown in Fig. 3B likely resulted from a cytostatic rather than an apoptotic response ^37,38^. In contrast, treatment with DOX-SSHEL^αHER2^ (carrying an equivalent of just 120 nM doxorubicin) induced caspase activity in SKOV3 after 24 h.

### DOX-SSHEL^αHER2^ particles enhanced HER2+ tumor clearance in a zebrafish brain metastasis model

The brain is often a site of relapsed metastatic disease for patients presenting with HER2^+^ breast cancers ^39^. We therefore sought to directly visualize the efficacy of SSHELs in clearing small tumor lesions in a complex tumor microenvironment using a zebrafish model of brain metastasis ^40^. We therefore co-injected ∼200 SKBR3 cells labeled with a fluorescent dye into the hindbrains of 3 days post-fertilization (dpf) zebrafish with various treatments. In zebrafish co-injected with PBS, SKBR3 cells migrated from the site of injection 5 h after injection and greater than 60% of the injected cells could be visualized within the brain 1 day after injection (Fig. 4A-A’; 4E-E’; 4I). When co-injected with ∼0.3 ng of doxorubicin or SSHEL^αHER2^, ∼60% of cells could still be detected near the site of injection 1 day after injection (Fig. 4B-B’; 4F-F’; 4I), indicating that the efficacy of doxorubicin was much less in vivo than it was in vitro (Fig. 3B). Co-injection of SKBR3 cells with DOX-SSHEL^αHER2^ at MOI-200 (equivalent dose of free doxorubicin) resulted in ∼60% tumor clearance within one day after injection (Fig. 4D-D’; 4H-H’; 4I): a statistically significant improvement compared to either treatment with PBS or doxorubicin. To test if potency depended on the size of the tumor, we plotted tumor clearance as a function of tumor size for individual zebrafish (Fig. 4J). SSHEL^αHER2^ (green) were most effective in clearance in relatively small tumor lesions (∼1-2 × 10^4^ square microns). In contrast, treatment with both DOX-SSHEL^αHER2^ (pink) and doxorubicin (blue) showed efficacy across a similar range of tumor sizes (0.1-6 × 10^4^ square microns), indicating that the combined action of cell targeting and drug delivery by DOX-SSHEL^αHER2^ resulted in increased tumor clearance in a zebrafish brain metastasis model, as compared to treatment with drug alone.

**Figure 4.**
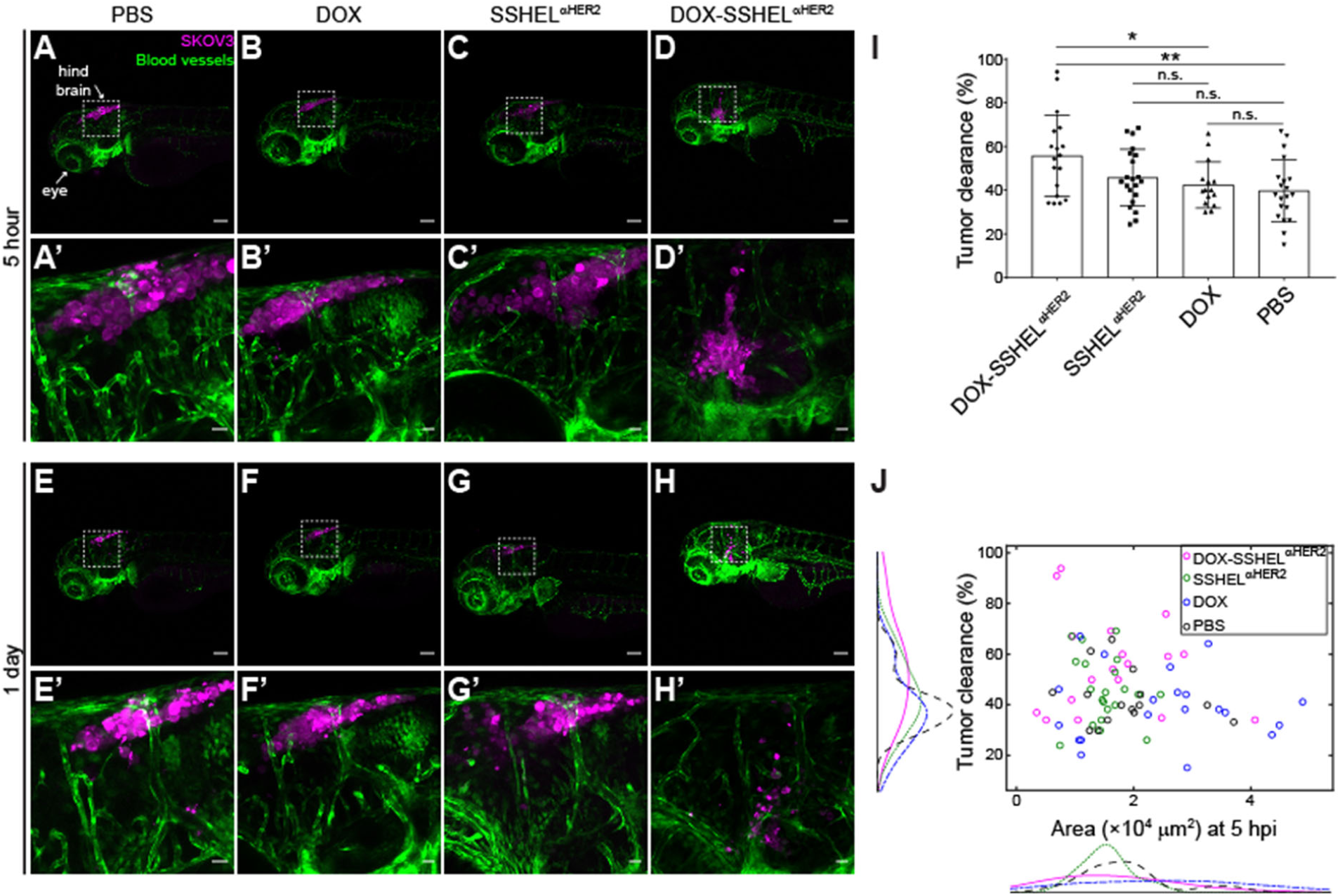
Doxorubicin-loaded SSHEL^αHER2^ enhances tumor clearance in a zebrafish brain metastasis model. (A-H’) Confocal micrographs of representative fli:GFP zebrafish (in which blood vessels produce GFP) 3 days post-fertilization were co-injected in the hind brain with ∼200 fluorescently labeled SKBR3 cells and (A) PBS, (B) 0.3 ng doxorubicin, (C) SSHEL^αHER2^ without doxorubicin, or (D) doxorubicin-loaded SSHEL^αHER2^ at MOI=200 (equivalent to 0.3 ng doxorubicin) imaged 5 h (A-D) and 1 day (E-H) after co-injection. (A’-H’) Magnification of square area indicated in (A-H), respectively. Images are overlays of fluorescence from GFP (green) and labeled SKBR3 cells (pink). Scale bar: 100 µm. (I) Quantification of tumor clearance measured using fluorescence microscopy, 1 day after co-injection with indicated treatment. Each data point represents tumor clearance in a single fish; bars represent mean; error bars are S.D. P-values: *<.05; **<.01; “n.s.”, not statistically significant. (J) Tumor clearance reported as a function of tumor size, for fish treated with PBS (black, n=15 fish), doxorubicin (blue, n=20), SSHEL^αHER2^ without doxorubicin (green, n=22), or doxorubicin-loaded SSHEL^αHER2^ (pink, n=19). Each data point represents tumor clearance in a single fish; histograms represent the aggregate of all the individual data along that axis.

## DISCUSSION

The efficacy of targeting antibodies to HER2 is thought to be mediated by three possible mechanisms: 1) inhibition of HER2 extracellular domain shedding that leads to internal degradation of the receptor; 2) reducing the number of binding sites for signaling ligands; and 3) stimulation of immune-regulated antibody-dependent cellular cytotoxicity in breast cancer ^23,41^. Trastuzumab (Herceptin) is thought function via the first mechanism, by successive degradation via ubiquitin ligase c-CBL via an unclear mechanism ^42^. In contrast, SSHELs bound to HER2 on the surface were internalized by the target cells. Since HER2 is not known to mediate endocytosis ^43^, we presume that the internalization occurred via a non-selective uptake mechanism such as macropinocytosis ^44^ or relatively slow endocytosis due to the relatively large size of the particle decorated with thousands of cell binding ligands, whereupon trafficking to an acidic compartment led to release of the SSHEL-encapsulated cargo and, in the case of a membrane-permeable drug like doxorubicin, delivery of the cargo to the cytosol. This mechanism is similar to that employed by Trastuzumab emtansine, an antibody drug conjugate (ADC) where the monoclonal antibody is covalently linked to DM1, a potent microtubule inhibitory drug that has been shown to be more potent than Trastuzumab alone ^23,45,46^. Our findings suggest that we can merge the benefits of this adjuvant therapeutic with the efficacy of the first in line treatment options and raises the possible beneficial effects of bystander killing where membrane-permeable drug is released from a killed cell.

Our observation that uptake results in potent cytotoxicity of both ovarian and breast cancer cells suggests a more generalizable strategy for targeting other cancers that have been less successful using current antibody base strategies. Although early clinical trials with Trastuzumab did not result in reduction of tumor burden in HER2^+^ ovarian cancer ^47,48^, in vitro and preclinical studies suggest that the drug may sensitize tumor cells to successive treatments against EGFR. It is therefore possible that SSHELs exploit a similar mechanism wherein the anti-HER2 affibodies affixed to the surface of SSHELs “primes” the target to be more susceptible to the encapsulated doxorubicin.

Myriad strategies employing nanoparticles and even live bacteria as drug delivery vehicles have been proposed for the specific killing of cancerous cells ^49-52^. We suggest that SSHELs provide several unique benefits and that they may be used as an additional strategy for this purpose and may be modified easily to suit different needs. First, SSHELs are composed of defined and biologically compatible materials. Second, SSHELs may be assembled in a straightforward and stepwise manufacturing process. Third, the discovery that low pH triggers the release of SSHEL-encapsulated cargo and the observation that they are internalized by the target cell and trafficked to an acidic compartment provides a specific cue that ensures the release of cargo at the correct place and time. Furthermore, the porous silica bead can accept a wide range of molecules, including proteins and nucleic acids, to vary the type of therapeutic delivered ^53^. Fourth, SSHELs exhibited low cytotoxicity, with mice tolerating multiple injections without any obvious negative response ^21^. Finally, the ability to covalently link (potentially multiple ^9^) ligands at high valency to the surface of SSHELs permits the active targeting of specific cell types, instead of passive diffusion to a tumor site due to vascular leakage. This is an attractive feature as de novo resistance to antibody-based treatment is thought to occur via compensatory mechanisms mediated by activation of other RTKs. Targeted therapies at the time of diagnosis have improved survival in HER2 breast cancer, however limited options are available for patients that show relapse in the brain. One explanation for this limited efficacy is that organ-specific cues found in the brain are distinct from conditions elsewhere examined in preclinical studies. A second possibility is reduced antibody penetrance. Our results demonstrated that SSHELs alone showed potency in the zebrafish brain microenvironment for small tumors and that cargo-loaded particles were more effective in the larger tumors, thereby further highlighting the benefit of combination therapy. We therefore envision that the platform we have presented here represents a versatile and safe “microparticle” whose targeting specificity and cargo may be modified to suit a specific application.

## MATERIALS AND METHODS

### Cell lines and culture

Human breast cancer cell lines (SKBR3, MDA-MB-231, MCF7) and human ovarian cancer cells (SKOV3) were purchased from the American Type Culture Collection. Cells were cultured in complete growth medium, consisting of RPMI 1640 medium (Gibco) supplemented with 10% fetal bovine serum (FBS, Gibco) and 1X penicillin-streptomycin (Gibco). All cells were incubated at 37 °C in 5% CO_2_.

### Protein purification and labeling

His_6_-SpoIVA harboring a single engineered N-terminal Cys was overproduced in *E. coli* “*ClearColi*” BL21(DE3) (Lucigen), which harbors a modified LPS that does not trigger endotoxic response, and purified using Ni^+2^ affinity chromatography and ion-exchange chromatography as described previously ^54^. For click chemistry conjugation, SpoIVA was labelled with 10-fold molar excess of *trans*-cyclooctene-PEG3-Maleimide (Click Chemistry Tools) overnight at 4 °C, after which the excess reagent was removed by Zeba Spin Desalting Column (Thermo Scientific). Anti-ErbB2 affibody molecule (Ab31889) was purchased from Abcam and labelled with Methyltetrazine-PEG4-maleimide. For labeling Ab31889 with fluorophore Alexa 647, affibody was incubated with 3-fold molar excess of Alexa 647 Succinimidyl Ester (Life Technologies) for 1 h at room temperature and desalted with Zeba Spin Desalting Column prior to labeling with Methyltetrazine.

### SSLB preparation

Liposomes were produced by the liquid sonication method using 200 µl (25 mg/ml) of 1,2-dioleoyl-*sn*-glycero-3-phosphocholine (DOPC, Avanti), that were first evaporated under vacuum for 3 h at room temperature and hydrated in 1 ml 0.5X PBS as described previously ^33,55^. Briefly, resuspended lipids were subjected to five freeze-thaw cycles between ethanol-dry ice bath and 42 °C water bath, followed by bath sonication until the suspension became transparent. Debris was removed by centrifugation at 13,000 × g for 10 min, and the supernatant containing small unilamellar vesicles was retained. One micron-diameter mesoporous silica beads (pore size of 10 nm) (Glantreo) were prepared for coating by washing three times each in 1 ml ultrapure water, followed by methanol and 1M NaOH. Beads were then rinsed and resuspended in ultrapure water (12.5 mg/ml). SSLBs were constructed by mixing 200 µl beads with 200 µl prepared liposomes at room temperature for 90 min with occasional pipetting. SSLBs were collected by centrifugation at 13,000 x g for 2 min, washed three times with 0.5X PBS and resuspended in 200 µl 0.5 X PBS.

### SSHEL assembly

SSHELs were made largely as described previously ^9^. Briefly, synthesized SpoVM peptide (Biomatik Corp.) was incubated at 10 μM (final concentration) with 2.5 mg/ml SSLBs in 0.5X PBS, at room temperature for 3 h with occasional pipetting. SpoVM-coated SSLBs (1 mg) were collected by centrifugation at 13,000 × g for 2 min and incubated with ∼1.5 µM *trans*-cyclooctene-labeled SpoIVA in a final volume of 400 μl PBS containing 10 mM MgCl_2_ and 4 mM ATP, overnight at room temperature with gentle inversion in a darkened test tube. To make SSHEL^αHER2^, SSHELs were collected by centrifugation, resuspended in 400 µl PBS containing 1.5 µM methyltetrazine-labeled Ab31889, and incubated overnight at room temperature with gentle inversion. SSHEL^αHER2^ were collected by centrifugation, washed thrice with PBS, and stored in PBS at 4 °C. Fluorescence microscopy images of SSHELs were obtained using Delta Vision Core microscope with a Photometrics CoolSnap HQ2 camera (Applied Precision) as described previously ^56^. For flow cytometry and confocal microscopy assays, 2 mg SSHELs were labeled with 10 µM of Alexa 488 Succinimidyl Ester (Life Technologies) for 1 h at room temperature prior to conjugation with affibodies. To estimate the copy number of Ab31889 conjugated onto SSHELs, the degree of labeling of Alexa 488 to Ab31889 was first estimated using absorbance measurements at 280 nm and 495 nm, and a correction factor of 0.11 for the AlexaFluor488 dye (Molecular Probes). The median fluorescence intensity (MFI) of SSHEL particles containing AlexaFluor488-labeled Ab31889 was measured using flow cytometry (FACSCanto II, BD Biosciences) and quantified using BD FACSDiva software. The MFIs were converted to molecules of equivalent soluble fluorochrome (MESF) of the SSHELs via a standard curve relating the fluorescence peaks of commercial Quantum™ AlexaFluor488 beads to their MESF values (Bangs Laboratories, Inc.); the estimated fluorophore to protein ratio (∼0.25) was used for calculating the Ab31889 copy number.

### Drug loading

Mesoporous silica beads (25 mg) were soaked in 200 µl doxorubicin hydrochloride (5 mM in water, Sigma) for 1 h at room temperature. Excess drug was removed via centrifugation at 13,000 × g for 1 min and discarding the supernatant. After loading, targeted SSHELs were constructed as described above. To determine the capacity of SSHELs for doxorubicin, 1 × 10^8^ doxorubicin-loaded SSHEL^αHER2^ were incubated in 1 % SDS (dissolved in 1 ml PBS) for 24 h with inverting at 37 °C. After centrifugation (13,000 x g, 2 min), the fluorescence intensity of doxorubicin (excitation, 470 nm; emission, 585 nm) in the supernatant was measured using a fluorescence microplate reader (SPARK, Tecan) and the concentration of doxorubicin in the supernatant was calculated using a standard curve.

### Cell binding assay

Cells grown in T-75 flasks (Corning) to 70-80% confluence were harvested with 10 mM EDTA in PBS for 15 min at 37 °C. Cells were washed twice with Iscove’s Modified Dulbecco’s Medium (IMDM, Gibco) supplemented with 10% FBS by centrifugation at 1,000 × g, for 2 min and resuspended in IMDM/FBS. Increasing ratios (multiplicities of incubation ranging from 20 to 150) of SSHEL^αHER2^ (labeled with AlexaFluor647) were incubated with 5 × 10^5^ cells/ml in siliconized tubes (G-tubes, Bio Plas Inc.) for 1 h at 37 °C with gentle inversion. Cells were then washed with PBS once, resuspended in 1 ml PBS, and examined by flow cytometry (FACSCanto II, BD Biosciences). All data analysis was performed using FlowJo software (Tree Star). For confocal live cell imaging, 2.5 × 10^5^ cells were seeded in poly-d-lysine-coated 35 mm glass-bottom dishes (MatTek) in 2 ml complete growth medium. After 24 h, cells were co-incubated with 5 x 10^7^ SSHELs ^αHER2^ labeled with AlexaFluor488 for 1 h at 37 °C. Cells were then washed twice with PBS, once with OptiMEM (Gibco), and stained with Hoechst 33342 (1 μg/ml in OptiMEM) for 30 min at 37 °C. Samples were washed three times with PBS and visualized using confocal microscopy (Zeiss LSM780).

### Cell viability and apoptosis assays

Cell viability was assessed by using cell counting kit-8 (CCK-8, Dojindo Molecular Technologies, Inc.) according to the manufacturer’s instructions. Briefly, 100 μl of EDTA-detached cells (50,000 cells/ml complete growth medium) were seeded in 96-well plates (Corning) and allowed to grow for 24 h. Cells were treated with 100 μl of complete medium containing either various forms of SSHELs, free doxorubicin, or affibody alone, and incubated for 90 h at 37 °C. The culture medium was removed and replaced with IMDM containing 10% (v/v) CCK-8 solution and further incubated for 2.5 h at 37 °C. Absorbance at 450 nm was measured by using multi-well plate reader (SpectraMax 190, Molecular Devices). Cell viability was calculated as a percentage of non-SSHEL treated sample. Caspase-3/7 activities were measured using Caspase-Glo 3/7 Assay kits (Promega) according to manufacturer’s instructions. Briefly, cells were seeded in 100 μl of complete medium at a density of 5,000 cells/well in a white 96-well plate (PerkinElmer). After 24 h incubation, cells were treated with 100 µl of complete medium containing either free doxorubicin (240 nM), doxorubicin-loaded SSHELs^αHER2^ (equivalent dose of free doxorubicin), or cargo-free SSHELs^αHER2^. Culture medium was removed at various time points and replaced with PBS containing 50% (v/v) Caspase-Glo reagent. The sample plate was incubated for 30 min at room temperature in darkness and read on a Victor3 luminometer (PerkinElmer).

### Confocal microscopy

SKOV3 cells were plated on 35 mm glass-bottom dishes (MatTek) at a density of 2 × 10^5^ cells in 2 ml complete growth medium and incubated for 24 h. Cells were co-incubated with 5 × 10^7^ Alexa 488-labeled SSHELs ^αHER2^ for indicated time periods, washed twice with PBS, once with OptiMEM (Gibco), and stained with Hoechst 33342 (1 μg/ml in OptiMEM) for 30 min at 37 °C. After washing once with OptiMEM, the acidic organelles of cells were stained with 300 nM LysoTracker Red DND-99 (Invitrogen) for 30 min at 37 °C. Cells were washed three times with PBS and visualized under a confocal microscope (Zeiss LSM780).

### In vitro drug release assay

To evaluate the doxorubicin release kinetics, 1 × 10^8^ doxorubicin-loaded SSHELs^αHER2^ were resuspended in 1 ml of physiological buffer (PBS with 10% FBS, pH 7.4) or 0.1 M citrate buffer (pH 5.0) and incubated at 37 °C with inverting. At defined time points, particles were removed via centrifugation (13,000 × g, 2 min) and fluorescence intensity of the supernatant was measured using a microplate reader. The concentration of released doxorubicin was converted into a percentage of the doxorubicin that was initially encapsulated within 10^8^ particles.

### Animal studies

Animal studies were conducted under protocols approved by the National Cancer Institute, and the National Institutes of Health Animal Care and Use Committee.

### Mouse experiments

Athymic nude mice (4-6 weeks old) were subcutaneously implanted with 1 x 10^7^ SKOV3 cells in 100 µl of 1:1 PBS/Matrigel (Corning) in the right flank. Tumor volume was determined by caliper measurement with the formula (V = length x width^2^)/2) and expressed as mean ± standard deviation. When tumor volume reached ∼100 mm^3^, mice were randomly divided into four groups (n=7 mice in each group): (1) vehicle control (PBS), (2) free doxorubivin (3 mg/kg), (3) DOX-SSHEL^αHER2^, and (4) empty SSHEL^αHER2^. Mice were injected via tail vein twice a week for a total of 11 injections. Tumor volumes and body weights were monitored throughout the treatment. Statistical analysis was performed by using the nonparametric Kruskal-Wallis test, with p < 0.05 being considered significant.

### Zebrafish

We employed the transgenic line, Tg(kdrl:GFP)la116, kindly provided by Brant Weinstein (NICHD, Bethesda, MD). Zebrafish were maintained at 28.5°C on a 14-hour light/10-hour dark cycle according to standard procedures. Larvae obtained from natural spawning, raised at 28.5°C and maintained in egg water containing 0.6 g sea salt per liter of DI water were checked for normal development. At day 5 regular feeding commenced. For all experiments, at 24 hours post fertilization (hpf), larvae were transferred to egg water supplemented with phenylthiourea (PTU, Sigma P5272), suspended at 7.5% w/v in DMSO, at 1 part in 4500 to inhibit melanin formation for increased optical transparency. Larvae were then returned to the incubator at 28.5°C and checked for normal development. Zebrafish larvae at 2 days post fertilization (2 dpf) were anesthetized using 0.4% buffered tricaine. Human cells were labeled with cell tracker (Deep Red, Invitrogen) at a final concentration of 1 µM for 30 minutes as previously described ^57,58^. Cells were then centrifuged, supernatant was discarded, and the pellet was resuspended in PBS with a final cell concentration of 1 × 10^7^ cells/mL before injection. 2-5 nL of labeled cells were co-injected into the hindbrain parenchyma of the larvae with PBS, 0.3 ng doxorubicin, SSHEL^αHER2^ without doxorubicin at MOI=200, or doxorubicin-loaded SSHEL^αHER2^ at MOI=200 (equivalent to 0.3 ng doxorubicin). Zebrafish were then reared at 28.5°C with 5% CO_2_ and relative humidity maintained at 95% for two days. At 5 days post fertilization, some zebrafish were then returned to system water and regular feeding at 28.5 °C for long-term survival studies. Zebrafish were screened within 24 h of injection to check for successful introduction of cells.

For live cell imaging, larvae were anesthetized using 0.4% buffered tricaine, then embedded in a lateral orientation in 1% low melting point agarose (NuSieve GTG agarose, Lonza), and allowed to polymerize in with cover glass (no. 1.5 thickness) as previously described ^59^. Egg water supplemented with tricaine was added to the agarose hydrogel for the entire time of data acquisition. Zebrafish were imaged on Zeiss 710 or 780 laser scanning confocal microscopes. Initial overview experiments were taken at 20×, with 1 µm Z steps, as tile scans over the entire length and height of the zebrafish. Images in the head and tail of the zebrafish were acquired at 10× magnification every 10 min for 14 h. Z stacks were acquired using a tiled approach and a 10× air objective of 0.3 NA where each individual image comprised 2046 × 2046 square pixels corresponding to 1416 × 1416 square µm for a total Z distance of 276 µm. One-photon, confocal, 2-dimensional images of 512 × 512 pixels (lateral dimensions) were acquired with a 1.4 NA 40 × oil-immersion objective. We simultaneously excited our sample with the 405 nm, 488 nm lines from an argon ion laser with a power of < 3 % (total power 30 mW) and 546 nm from a solid-state laser (power of < 10 %). A secondary dichroic mirror, SDM 560, was employed in the emission pathway to delineate the red (band-pass filters 560–575 nm) and green (band-pass filters 505–525 nm) and blue channels (480-495 nm) at a gain of 400 on the amplifier. The laser power for the 543 nm setting was set at < 3 % of the maximum power and the gain on the detectors was set to 450.

### Quantification of in vivo tumor clearance-Zebrafish

Image processing was performed via ImageJ. Spectral crosstalk was removed by subtracting contributions of the red channel from the far-red detection channel. Images were then median (radius of 3 pixels) and Gaussian (sigma of 2) filtered to minimize background noise. Binary masks of cells were generated using Huang and Otsu thresholding methods on corrected images. The area occupied by cells was then calculated using ImageJ’s built in ‘analyze particles’ function. Two-way ANOVA was used to determine statistical significance.

## ACKNOWLEDGEMENTS

We thank members of our labs for discussion and comments on the manuscript. This work was funded by the Intramural Research Program of the National Institutes of Health, National Cancer Institute, Center for Cancer Research and by a Center for Cancer Research FLEX Synergy Award (to K.S.R., K.T., and D.J.F.).

## AUTHOR CONTRIBUTIONS

M.K., D.J.F., K.T., and K.S.R. designed research; M.K., T.U., M.A.C., D.G., and I.W. performed research; M.K., M.A.C., D.J.F., K.T., and K.S.R. analyzed data; M.K., D.J.F., K.T., and K.S.R. wrote the paper.

---

**Figure S1.**
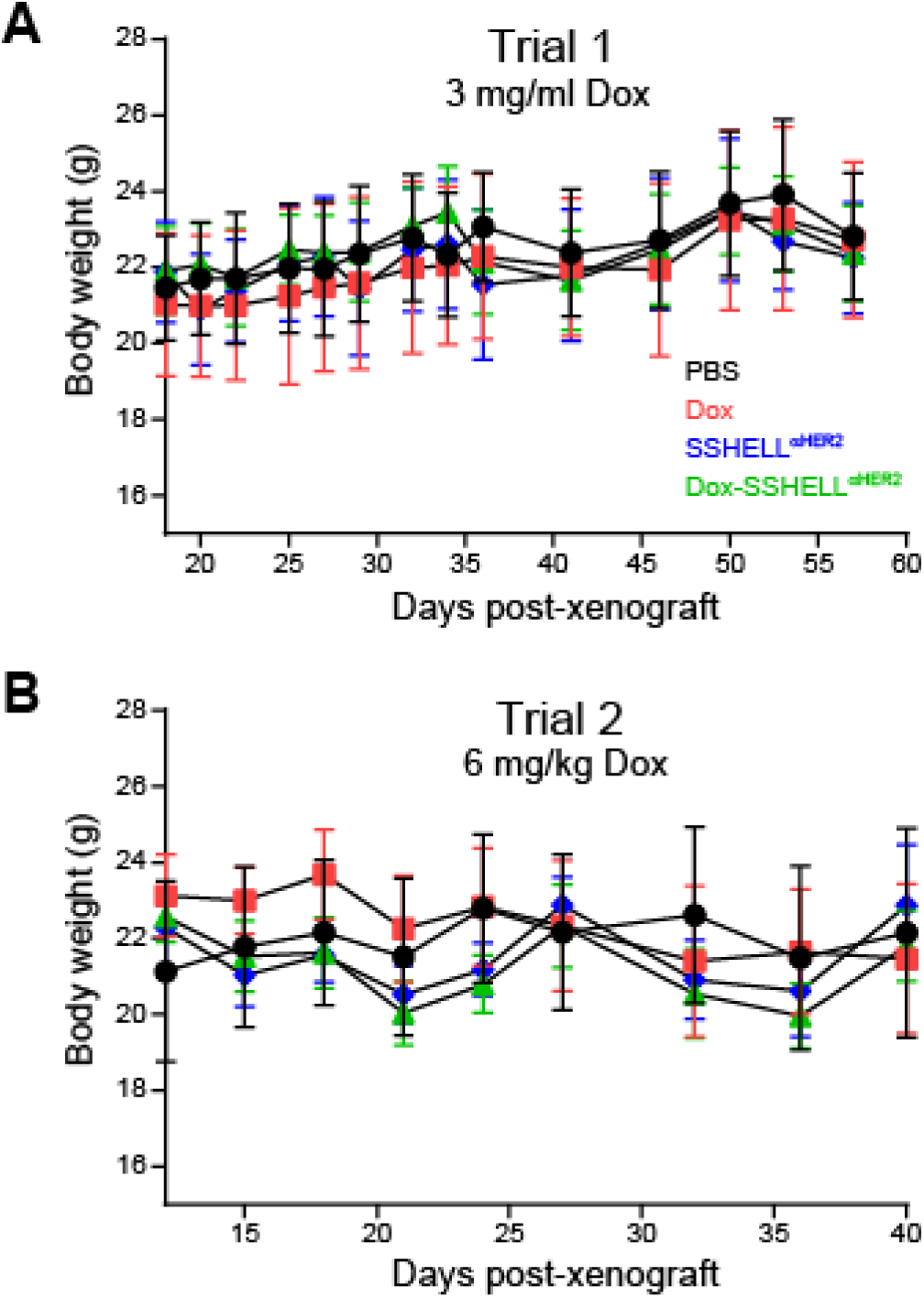
Average body weight of athymic nude mice harboring SKOV3 tumor xenograft. Mice were weighed periodically after treatment began with PBS (black), free doxorubicin (red), unloaded SSHEL^αHER2^ (blue), or doxorubicin-loaded SSHEL^αHER2^ (green). (A) Trial 1 in which 3 mg/kg doxorubicin was administered; (B) Trial 2 in which 6 mg/kg doxorubicin was administered.

**Figure S2.**
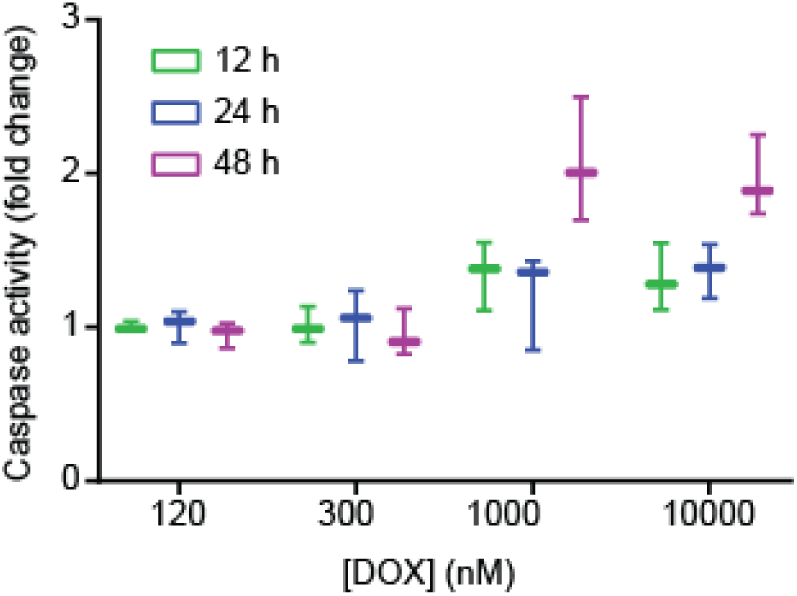
Modest induction of caspase activity in SKOV3 cells by doxorubicin. Caspase 3 and 7 activities of SKOV3 cells were measured using indicated concentrations of doxorubicin after 12 h (green), 24 h (blue), and 48 h (magenta) of treatment. Data represent the average fold change compared to PBS-treated cells; errors are S.D. (n=3).

